# Astrocyte regulatory volume decrease is condition-dependent in intact brain tissue and requires the volume regulated anion channel

**DOI:** 10.64898/2026.08.14.737967

**Authors:** S. Sriram, C. D. Lopez, P. Pham, D. K. Binder, T. A. Fiacco

## Abstract

Multiple lines of evidence point to the volume regulated anion channel (VRAC) as being instrumental for cellular volume regulation in many cell types, including astrocytes. VRAC are thought to open during periods of astrocyte swelling, releasing anions and osmolytes to drive water out of the cell, allowing it to return to baseline volume even under sustained osmotic or ionic challenge, a process called regulatory volume decrease, or RVD. However, the occurrence of RVD and VRAC’s role in this process has remained controversial, with clear evidence in cultured cells but mixed reports from work in intact brain tissue. In the present study, we aimed to address this gap by generating a transgenic mouse line in which VRAC is conditionally ablated in astrocytes (VRAC cKO) and recording the volume responses of astrocytes in VRAC cKO and control tissue using real-time volume imaging. We found that the effect of VRAC cKO on astrocyte swelling was dependent on whether swelling was evoked by elevated extracellular potassium, or by reduced extracellular osmolarity. We also found that both VRAC and the presence of sufficient intracellular taurine concentration were required to elicit RVD in astrocytes, but only in hypoosmolar conditions. Our findings provide new information on the conditions needed to elicit RVD in intact brain tissue, and that VRAC is required for RVD to occur. Our findings further suggest that reduction of intracellular ion concentration is essential for VRAC to be activated, rather than simply membrane expansion. Future experiments will examine the solute release aspect of VRAC activation upon astrocyte swelling, as well as the contributions of VRAC to pathological volume dysregulation.

## Introduction

The volume and composition of brain’s extracellular space (ECS) is critical for healthy neuronal function. ECS volume is inversely proportional to cellular volume, which expands and contracts in response to physiological and pathological changes in neuronal activity (Fultz et al., 2019; Sriram et al., 2024; Xie et al., 2013). Accumulating evidence suggests that astrocytes play a key role in ECS composition and volume due to their primary role in ion and neurotransmitter update (Herminia Pasantes-Morales & Vázquez-Juárez, 2012). Medical conditions featuring fluid dysregulation have been well documented, their causes ranging from structural perturbation following physical injury to disordered fluid intake behaviors (Hannon et al., 2012; Murphy et al., 2017). The resulting ionic imbalances can disrupt astrocytic maintenance of homeostasis, potentially causing or exacerbating damage to the brain as part of a long-term pathology response (Dreier et al., 2018).

It is generally understood that all animal cells are equipped with compensatory mechanisms against sustained swelling such that, upon exposure to osmotic or ionic challenge, the initial swelling response is followed by an attempt to return to baseline volume (Mongin & Orlov, 2001). This process, known as regulatory volume decrease (RVD), has been studied extensively in a variety of contexts, including epithelial cells, lymphocytes, and nephrons (Cheung et al., 1982; Kirk et al., 1987; Okada et al., 2001). In the same way extracellular solute imbalance leads to water entry into cells, efflux of ions is the predominant driving force for water extrusion, enabling the cell to return to baseline volume (Pasantes-Morales et al., 1994; Sarfaraz & Fraser, 1999). Specifically, swollen cells have increased anion permeability and display an outwardly rectifying chloride current (Cahalan & Lewis, 1988; Grinstein et al., 1982; Hazama & Okada, 1988; Olson & Li, 1997). In the brain, this outward anionic current is typically accompanied by efflux of neuroactive osmolytes, like glutamate and taurine, the release of which is implicated in several pathological states (Abdullaev et al., 2006; L. Chen et al., 2019; Kimelberg et al., 2004; Ramírez-Guerrero et al., 2022).

Overall, multiple lines of evidence suggest that activation of the swelling-activated chloride current is indispensable for RVD (Grinstein et al., 1982; Hazama & Okada, 1988). Early investigations into a potential underlying channel discovered the volume-sensitive organic anion channel (VSOAC), but its specific molecular identity was unknown for many years (Hoffmann et al., 1984; Jackson & Strange, 1995; Okada, 1997). Following attempts by several labs to elucidate the mechanism, the Jentsch and the Patapoutain groups discovered that the LRRC8 family of proteins are essential for activation of this current (Qiu et al., 2014; Voss et al., 2014). VRAC is a hexameric transmembrane channel composed of LRRC8 subunits, LRRC8A-E, each encoded by genes *LRRC8A-E.* VRAC channels may be composed of any combination of LRRC8 subunits, and a functional channel must contain at least one LRRC8A subunit (also referred to as SWELL1) (Voss et al., 2014). VRAC is permeable to several anions and organic osmolytes following an Eisenmann type 1 permeability profile: SCN^−^ > I^−^ > NO_3_^−^ > Br^−^ > Cl^−^ > formate > propionate = methanesulphonate = acetate ≥ F^−^≥ butyrate > valerate > gluconate = glucuronate = glutamate (Ghouli et al., 2022; Nilius & Droogmans, 2003). VRAC activates upon cell volume increase, triggering an efflux of anions, namely chloride, and organic osmolytes, like glutamate, aspartate, and taurine (Feustel et al., 2004; Formaggio et al., 2019; Qiu et al., 2014).

While RVD has been historically observed in a variety of cell types (Okada et al., 2001), there is conflicting evidence regarding its occurrence and mechanisms. This is likely due to the parameters that vary drastically across experimental models such as cell type, swelling trigger, time course, pharmacology, recording temperature, imaging techniques, and tissue preparation methods (Haskew-Layton et al., 2008). For example, while cultured peripheral lymphocytes show a robust RVD in response to hypotonic challenge, red blood cells require hours to return to baseline volume (Cheung et al., 1982). In the brain, Pasantes-Morales et al. (1994) showed robust and dramatic compensatory volume changes within 6 minutes of exposure to hypoosmotic conditions, while Andrew et al. (1997) provided evidence against volume regulation by cortical brain cells during acute osmotic stress. Even among those that have observed RVD, it remains controversial which specific astrocytic channels participate in compensatory volume changes beyond *in vitro* preparations (Lafrenaye & Simard, 2019; Mola et al., 2021; Walch & Fiacco, 2022).

Astrocytic VRAC has been heavily implicated in the anion and osmolyte efflux associated with water expulsion and compensatory volume changes. However, methodological constraints have made it difficult to determine the contribution of VRAC to astrocytic volume changes in intact tissue. Our lab has previously used real-time volume imaging in acute hippocampal tissue to demonstrate neuronal and astrocyte swelling in hypoosmolar conditions and astrocyte-selective swelling in elevated extracellular potassium ([K^+^]_o_ (Walch et al., 2020). In this study, we applied these techniques to explore the conditions and contexts for RVD in hippocampal CA1 stratum radiatum astrocytes. We also sought to establish the contribution of astrocytic VRAC to volume regulation using a transgenic strategy to ablate LRRC8A selectively in astrocytes. Here we show that both the occurrence of RVD and the role of VRAC in cell volume regulation is condition-dependent. RVD occurred in hypoosmolar conditions only when near-physiological levels of taurine were present. In the absence of taurine, functional VRAC capped the maximal swelling response without evidence of RVD. Despite similar magnitude swelling responses, VRAC did not regulate astrocyte volume or generate RVD in elevated [K^+^]_o_ up to 26 mM. Last, RVD was completely absent in brain slices from VRAC cKO mice, identifying VRAC as a critical regulator of RVD and cell volume regulation in intact brain tissue.

## Materials and Methods

All experiments were performed in compliance with the National Institutes of Health guidelines for the care and use of laboratory animals, and protocols were approved by the Institutional Animal Care and Use Committee at the University of California, Riverside.

### Transgenic Mouse Breeding Strategy

*LRRC8A*, alternatively known as *SWELL1*, encodes for a subunit of the VRAC channel that is essential for the formation of the pore and for trafficking the channel to the membrane (Jentsch et al., 2016; Voss et al., 2014). We used the Cre-lox system to selectively target and ablate the *LRRC8A* gene in two complementary transgenic mouse lines to generate astrocyte-specific conditional knockouts (cKO). Double floxed LRRC8A (*SWELL^fl/fl^*) bred on a C57/Bl6 background were obtained from Dr. Rajan Sah at Washington University. These mice were crossed with a carrier of Cre recombinase driven by the mGFAP promoter, again on a C57/Bl6 background (Jackson Laboratories). Offspring carrying Cre recombinase and one allelic copy of floxed LRRC8A were bred back to SWELL^fl/fl^ mice until double floxed mice carrying Cre recombinase were produced (*mGFAP-Cre^+/-^; SWELL^fl/fl^*), hereby referred to as astrocyte VRAC cKO. Cre-negative double floxed mice (*mGFAP-Cre^-/-^; SWELL^fl/fl^*) from the same litters as knockout mice were used as controls.

### *In Situ* Hybridization and Analysis

We used fluorescent *in situ* hybridization to quantify LRRC8A mRNA in astrocyte VRAC cKO and littermate control mice. To prepare the tissue, animals were sacrificed at 8-12 weeks of age, perfused, and their brains harvested as described above. Following the overnight postfixation, brains were cryoprotected in 30% sucrose in 1X PBS for 1-2 days, or until they no longer floated in solution. They were then embedded in OCT (ThermoFisher) and sectioned sagittally at 20 µm on a cryostat (Leica CM1950). The slices were mounted onto polarized microscope slides and stored at -20° C for use within one month. We used the RNAscope Multiplex Fluorescent Reagent Kit v2 (Advanced Cell Diagnostics) following the manufacturer’s protocol (F. Wang et al., 2012). Additional reagents included diethyl pyrocarbonate (Millipore) to remove RNAses from assay solutions, a probe targeting *LRRC8A* (ACD 45837), and Opal Dye 520 (Akoya Biosciences) reconstituted in DMSO. Following the RNAscope assay, we then performed immunohistochemistry to visualize astrocyte morphology. Slides were blocked in 3% BSA in 1X PBS for 1 hour, then incubated with rat-anti GFAP with gentle agitation at 4° C overnight. Following several washes in 1X PBS, slices were incubated in AlexaFluor 488 and DAPI as described above, washed again in 1X PBS, cover slipped with mounting medium, sealed, and stored at 4° C until imaging.

For visualization and quantification, Z-stack images were acquired on a laser scanning confocal microscope (Leica SPEII) at 20X for *in situ* hybridization experiments to achieve high single-cell resolution of stratum radiatum astrocytes. When using higher magnification, images were taken in two directly adjacent areas of CA1 and quantification from both areas was combined for each slice. Max projections were created using LasX software (Leica), then imported into QuPath (Bankhead et al., 2017) for analysis. Regions of interest (ROIs) were drawn around the pyramidal layer and stratum radiatum of the hippocampus based on DAPI and GFAP density. We then used the “Cell Detection” function to identify astrocytes labeled with GFAP. To identify mRNA puncta exclusively within astrocytes, we used the “Subcellular Detection” function using GFAP signal to outline astrocyte soma and processes. Since GFAP labelling of astrocyte perivascular endfeet may lead to unwanted detection of endothelial *LRRC8A*, we manually removed cellular and subcellular detection around vasculature. Thresholds for minimum signal intensity and cell size were kept consistent for all images within an experiment.

To quantify astrocytic *LRRC8A* in stratum radiatum, total puncta area was normalized to the total amount of GFAP signal detected. To measure *LRRC8A* in CA1 pyramidal cells, astrocytic puncta area was measured as in the stratum radiatum, then subtracted that from the overall area of puncta in the stratum pyramidale. The area of the remaining pyramidal neuronal puncta was then normalized to the size of the pyramidal layer ROI. To account for loss of slices or damage to tissue during assays, quantification for each animal was based on data averaged from 2-4 slices.

### Acute Hippocampal Slice Preparation

Acute hippocampal slices were prepared from *LRRC8A* cKO mouse lines using the procedure outlined in (Walch et al., 2022), adjusted to optimize adult tissue viability. Animals were anesthetized by isoflurane inhalation, then transcardially perfused with ice cold slicing buffer bubbled with a mixture of 95% O_2_ and 5% CO_2_ and containing (in mM): 87 NaCl, 75 sucrose, 10 glucose, 1.25 NaH_2_PO_4_, 2.5 KCl, 25 NaHCO_3_, 1.3 ascorbic acid, 0.5 CaCl_2_, and 7 MgCl_2_, with an osmolarity of 315-325 mOsm. Following the perfusion, animals were immediately decapitated and brains rapidly and carefully harvested and placed in a partially frozen oxygenated slicing buffer (recipe above, with the addition of 2 pyruvate, 3.5 MOPS, and 0.1 kynurenic acid). While immersed in this solution, brains were sectioned on a vibratome (Leica VT1200S) into 350 µm thick parasagittal sections and trimmed until only the hippocampus and immediately adjacent cortex remained. These slices were then incubated in a chamber containing oxygenated slicing buffer which had been warmed 36° C in a hot water bath. The slicing buffer in the incubation chamber was supplemented with 4 µM sulforhodamine-101 (SR101, Sigma-Aldrich), which is selectively taken up by astrocytes (Schnell et al., 2012; Walch et al., 2020). After 45 minutes at 36° C, slices were brought to room temperature for 15 minutes, then transferred to a chamber containing oxygenated normal artificial cerebrospinal fluid (nACSF), which contained (in mM): 125 NaCl, 15 glucose, 1.25 NaH_2_PO_4,_ 2.5 KCl, 26 NaHCO_3_, 2.5 CaCl_2_, and 1.3 MgCl_2_, with an osmolarity of 300-305 mOsm. Slices remained here for 15 minutes before being placed in another chamber containing oxygenated nACSF at room temperature to allow for any residual SR101 to be washed from slices during the transfer. Slices recovered in the final chamber for a minimum of 30 minutes. Following recovery, slices were placed on top of a glass coverslip in a recording chamber (RC-26GLP, Warner Instruments) on the microscope stage and continuously perfused with freshly oxygenated solution. For experiments conducted at physiological temperature, the recording chamber was warmed to 36° C using a TC-344B Dual Automatic Temperature Controller system with an in-line solution heater set to 37° C (Warner Instruments). This slight discrepancy was intended to reduce the formation of gas bubbles upon heating, which may affect the perfusion rate of solutions as they passed to the recording chamber, as well as to account for any heat loss.

### Experimental Solutions and Pharmacology

All slices were first exposed to oxygenated nACSF upon placement into the recording chamber, and baseline volume for each cell was established with 3 z-stacks taken at 1-minute intervals. For experiments carried out at physiological temperature, slices were given 10 minutes to warm up to 36° C before acquisition of baseline images. To trigger astrocyte-specific swelling, [K^+^]_o_ was increased to either 10.5 mM or 26 mM K, maintaining osmolarity by removing a corresponding amount of NaCl (final concentration: 117 mM and 101.5 mM respectively). These values have been reported under moderate to severe pathological conditions (Heinemann & Lux, 1977). To evoke tissue-wide swelling, the osmolarity of ACSF was reduced by 40% (180-183 mOsm), either by lowering the concentration of NaCl or by diluting nACSF with diH_2_O (66.2 mM and 75 mM NaCl respectively). These modifications are collectively referred to as “experimental ACSF”.

For taurine experiments, following the procedure outlined in (Kreisman & Olson, 2003), the slicing buffer and ACSF recovery solutions used for incubation described above were supplemented with 1 mM taurine. Thus, slices were “loaded” with taurine for a minimum of 90 minutes before being transferred to the recording chamber. Once in the recording chamber, slices acclimated to physiological temperature for 10 minutes, during which they were continually exposed to 1 mM taurine ACSF. A 3-minute “pre-baseline” was established immediately after this time to examine any volume effects of basal taurine efflux alone (as opposed to swelling-evoked taurine efflux). Slices were then exposed to nACSF for 30 minutes, and a new baseline was established during the last 3 minutes of this period. Experimental ACSF was then applied for 40 minutes, followed by a 20-minute nACSF wash period as described above. Cells that did not respond within 10 minutes of the experimental period or did not exhibit immediate recovery upon wash of experimental solution were excluded from further analysis.

### Real-time Volume Imaging and Analysis

Real-time volume imaging was performed using an Olympus FluoView FV1000 confocal imaging system, using an Olympus LUMPlanFl 60X/0.90 W 1/0 water immersion objective lens, and Olympus FluoView 1000 software. SR101 was visualized with a 559 nm semiconductor laser and detected using a 624-724 nm bandpass filter. For all experiments (excluding laser light control experiments), laser output was kept at 1.5% to minimize fading and light-induced increases in cellular metabolism. Confocal aperture size, gain, and offset values were kept consistent between experiments, and the intensity was kept at a range of 750-900 to account for variability in SR101 loading between slices.

Real-time volume imaging was performed as previously described (Murphy et al., 2017; Walch et al., 2020). Briefly, slice health was assessed using differential interference contrast optics (Olympus), then SR101- labelled astrocytes in the s. radiatum were visualized and selected for imaging based on their characteristic, highly branched morphology, their depth within the tissue (≥25 µm), and high signal to background ratio. Low magnification images were taken to obtain a view of astrocytes to get an overall assessment of health and ensure no blebs or vasculature interfered with the selected cell. The imaging field was then cropped close to the cell soma to maximize resolution and reduce acquisition time to under 10 seconds, which was important for reducing potential laser effects and maximize temporal resolution. Only one cell was imaged per slice and, if more than two time points were missed, the cell was discarded. To accurately represent the entire 3-D structure of the cell, z-stacks were taken at 1-µm increments, and any drift caused by tissue swelling or solution flow was tightly monitored using rapid adjustments of the objective in the *x-*, *y-*, and *z-*planes between image acquisition.

Images acquired on FluoView software were then imported in to Fiji/Image J for analysis as previously published (Murphy et al., 2017; Walch et al., 2020, 2022). Briefly, z-stacks taken at each time point were aligned based on their widest, brightest point, then compressed into a single hyperstack, which was thresholded and binarized to remove background signal. The area of the cell soma was quantified and this value was used to approximate changes in cell volume, with the cell’s size at each time point represented as a percent increase or decrease relative to baseline. Representative images of cell volume were created by overlaying a pseudocolored magenta image at the end of an experimental period onto a green one from the beginning such that non-overlapping magenta areas represent swelling of the soma and green areas represent shrinking.

### Calculation of RVD

Qualitatively, a stereotypical RVD is characterized by an initial volume increase in response to a sudden ionic or osmotic trigger, after which cells begin to pump out fluid in an attempt to reverse swelling and return to baseline conditions (Pasantes-Morales et al., 1994). This turns into an “undershoot” as the challenge is alleviated, indicative of compensatory mechanisms that remain active both during and long after the threat has been removed (Andrew et al., 1997). We quantified this response based on the definition of RVD put forth by Kreisman and Olson (2003). Briefly, for each averaged dataset, we defined the volume at the “end of swelling” as the average of the last three time points in the experimental period to account for natural point-to-point variation in live-tissue recordings. We then calculated the “peak swelling” as the maximum volume achieved during the experimental period averaged with the time point directly before and directly after it. We used the ratio between these two values, expressed as a percentage, to assess the amount of recovery, or the lack thereof. If astrocytes reached their maximum value at the end of application of swelling-inducing ACSF, this ratio was close to 100% and RVD was considered not to occur. If this value was below 80%, we determined this to be indicative of RVD.

## Statistical Analyses

For RNAscope characterization, stratum radiatum LRRC8A puncta count was normalized to GFAP area and averaged across 2-4 slices for a given animal. Since some astrocyte processes and cell bodies are present in the pyramidal layer, we first quantified the astrocytic LRRC8A, then subtracted that from the total puncta area in the pyramidal layer before normalizing it to pyramidal layer area. To analyze LRRC8A puncta in VRAC cKO astrocytes and in the pyramidal layer, we first performed a Shapiro-Wilk test to verify these data were normally distributed. Then, because Levene’s test for equality of variance was violated, we used Welch’s one-tailed independent samples t-test to determine statistical significance.

For volume imaging experiments, differences in astrocyte swelling responses between genotypes were measured. Outliers were detected based on the interquartile range (IQR) of the distribution as demonstrated by a boxplot (Tukey’s method). Cell volume at a given time point was considered an "extreme outlier" if the distance from the edge of the box exceeded three times the IQR. If a cell contained extreme outliers at three or more time points, it was excluded from the analysis. Significance values reported in the “Results” section represent results of interactions between time exposed to experimental ACSF and genotype obtained from a two-way mixed ANOVA. The assumption of sphericity was violated for nearly all datasets, so the Greenhouse-Geisser estimate was used to interpret interactions. For datasets with statistically significant interactions between solution exposure and genotype, and a main effect of genotype, ANOVAs were followed up with Mann-Whitney U tests to assess differences between genotypes at individual time points. Significance values reflecting these differences are represented by asterisks within figure panels as follows – ∗p < 0.05, ∗∗p< 0.01, and ∗∗∗p < 0.001.

## Results

### Astrocytic LRRC8A expression is significantly downregulated in *mGFAP-Cre^+/-^; SWELL^fl/fl^* hippocampal tissue

The Cl^-^ channel inhibitors DCPIB and tamoxifen have been shown to robustly inhibit VRAC, but their nonspecific effects have made it difficult to confirm the contributions of astrocytic VRAC to ECS volume and composition (Bowens et al., 2013). Additionally, dramatic differences in mechanisms of swelling-induced osmolyte efflux arise when comparing *in-vivo* and *in-vitro* preparations (Haskew-Layton et al., 2008), necessitating the development of a model where astrocytic VRAC may be studied in a more physiologically relevant context. Among the various LRRC8 subunits that may compromise a single VRAC channel protein (Ghouli et al., 2022), LRRC8A is critical for the formation of the channel pore and for its trafficking to the cell membrane (Formaggio et al., 2019; Qiu et al., 2014; Voss et al., 2014). Therefore, we used the Cre-lox transgenic system to generate a mouse line with astrocyte-specific ablation of VRAC under the *mGFAP* promoter to target the *LRRC8A* gene in astrocytes. There was no obvious behavioral phenotype stemming from selective knockdown of VRAC in astrocytes, and mice exhibited typical breeding behavior and lived normal lifespans. In real-time volume imaging experiments, there were no observable differences in neuronal health, baseline starting astrocyte volume, SR101 loading, or astrocyte morphology between genotypes, suggesting that astrocytic VRAC cKO did not significantly impair slice viability or other intrinsic tissue properties.

We combined immunohistochemistry with fluorescent *in-situ* hybridization to verify the absence of astrocytic *LRRC8A* in adult (8- to 12-week-old) *mGFAP-Cre^+/-^; SWELL^fl/fl^* hippocampal tissue. We then used QuPath to detect LRRC8A-labeled puncta within GFAP-labelled astrocyte soma and processes. In control tissue, LRRC8A mRNA was abundant in stratum radiatum astrocyte soma and processes, as well as in neurons and astrocytes in the pyramidal layer (**Fig. 1A, C**). In VRAC cKO astrocytes, LRRC8A mRNA was virtually absent (**Fig. 1B, D**), while puncta were still visible in the parenchyma, likely belonging to other neuronal and glial cell types. Reduction of mRNA in astrocytes is quantified in **Fig 1E** (Welch’s t (5.56) = 2.95, p = 0.01). To further validate the specificity of our transgenic model, we examined LRRC8A expression in the pyramidal layer and found no significant reduction of mRNA (**Fig 1F**), suggesting that VRAC had been largely preserved in neurons (Welch’s t (9.49) = 0.877, p = 0.20).

**Figure 1:**
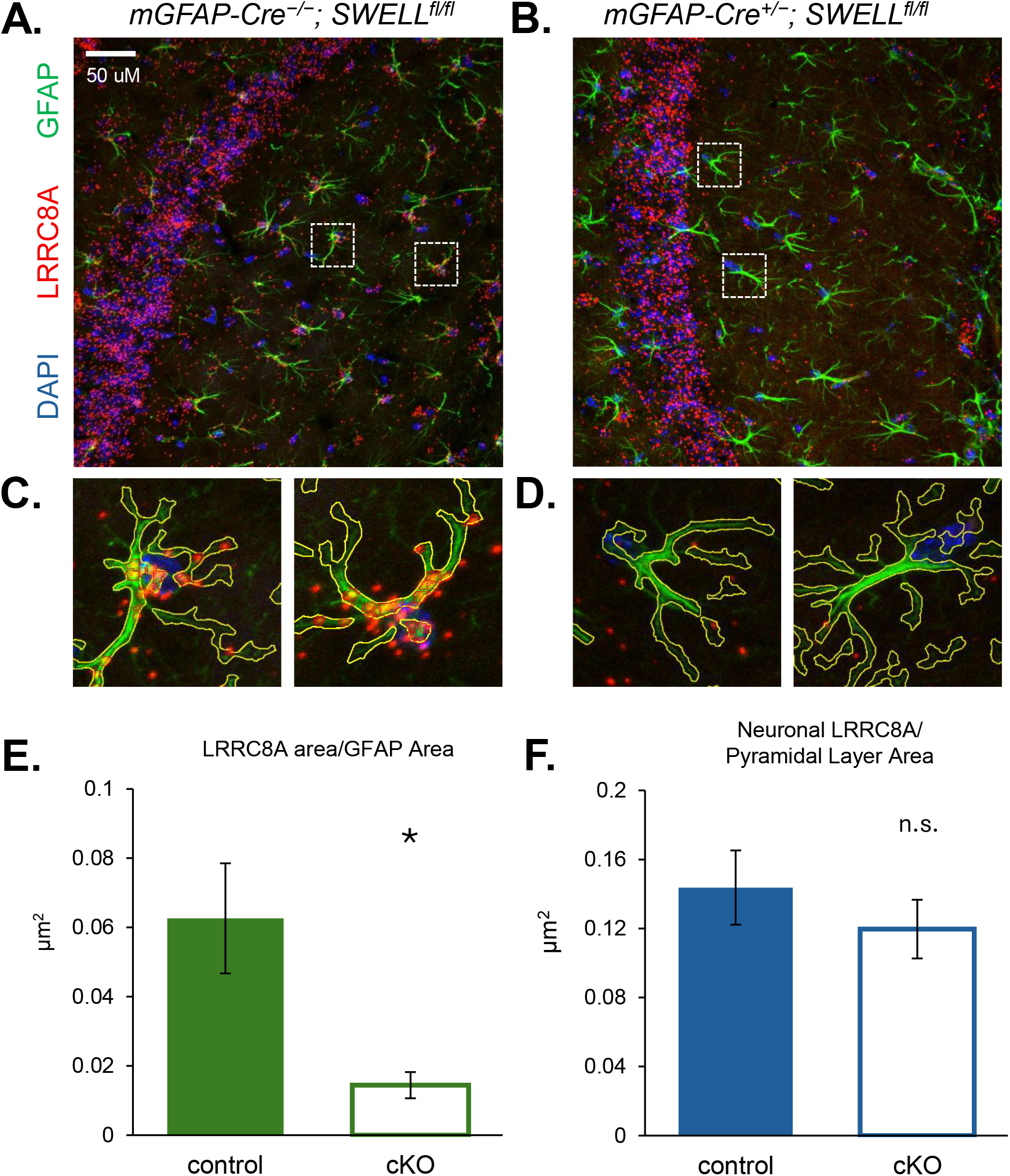
Validation of transgenic strategy for selective LRRC8A knockdown in astrocytes. Representative RNAscope/IHC images from control **(A)** and *mGFAP-Cre;SWELL ^fl/fl^* **(B)** astrocytes in the stratum radiatum of the hippocampus. **(C)** and **(D)**: Individual astrocytes from A and B (inset) showing cell morphology with GFAP (green) and LRRC8A puncta in red, detected using QuPath software. Note that astrocytes in control tissue have LRRC8A mRNA in both soma and processes, as indicated by red puncta surrounded by a red border, while VRAC signal is greatly reduced in *mGFAP-Cre;SWELL ^fl/fl^* astrocytes. **(E)** Quantification of LRRC8A area reveals that *mGFAP-Cre;SWELL ^fl/fl^* astrocytes have significantly less LRRC8A mRNA labeling compared to control astrocytes. **(F)** Abundant expression of LRRC8A mRNA remains in the pyramidal cell layer in *mGFAP-Cre;SWELL ^fl/fl^* sections, indicating selective knockdown of VRAC in astrocytes. *: p<0.05

### VRAC does not play a role in the astrocyte volume response to elevated [K^+^]_o_

We first set out to establish the contributions of VRAC to the long-term astrocytic volume response in 10.5 mM [K^+^]_o_. Our lab has previously demonstrated that, after 5 minutes of exposure to isoosmolar 10.5 mM [K^+^]_o_, astrocytes swell approximately 6.6% above their baseline volume, driven mainly by increased activity of the astrocyte Na^+^/K^+^ ATPase (Walch et al., 2020). We reasoned that VRAC may play a role in astrocytic volume regulation in these more physiological conditions of astrocyte swelling, which could occur during heightened and sustained neuronal activity. Due to the variable time course of observed RVD across several studies (Andrew et al., 1997; Okada et al., 2001; Pasantes-Morales et al., 1994), we extended the swelling protocol to 40 minutes of ionic challenge, followed by a 20-minute wash period.

The presence of ATP is thought to be critical for the swelling-induced anion current (Jackson et al., 1994; Rutledge et al., 1999) and the concurrent release of organic osmolytes (Hyzinski-García et al., 2014; Kimelberg, 2004; Mongin & Kimelberg, 2005). Both ATP and its poorly or non-hydrolysable analogues are thought to be necessary to potentiate swelling-induced anion current in cultured cells (Jackson et al., 1994). Extracellular ATP binds to P2Y receptors on astrocytes, triggering the release of intracellular Ca^2+^ stores, subsequently activating PKC and calmodulin pathways, both of which are thought to modulate activation of VRAC (Ghouli et al., 2022; Mongin & Kimelberg, 2005). Thus, 10.5 mM [K^+^]_o_ ACSF was supplemented with 10 µM Na-ATP to strengthen possible contributions of VRAC to any potential RVD (Kimelberg, 2004; Mongin & Kimelberg, 2005).

As described above, SR101-labeled astrocytes in stratum radiatum of CA1 hippocampus were selected based on their stereotypical shape and high signal-to noise ratio with distinct boundaries (**Fig. 2A, far left**). To visualize swelling and shrinking over the course of volume imaging, we used the thresholded and binarized z-stacks of astrocyte soma at various time points as described previously (Lauderdale et al., 2015; Walch et al., 2020). Astrocyte volume at baseline (**Fig. 2A, middle left)** was then pseudocolored green and overlaid with the z-stack taken 40 minutes after exposure to experimental ACSF, colored magenta (**Fig. 2A, middle right**). Magenta regions indicate where the membrane expanded relative to the baseline. To visualize the return to baseline upon removal of ionic or osmotic challenge, the 40-minute timeline was pseudocolored green and overlaid with the z-stack obtained at the end of the wash period, colored magenta (**Fig. 2A, far right**). As a result, green regions indicate recovery of cell volume relative to the point of maximum swelling.

**Figure 2:**
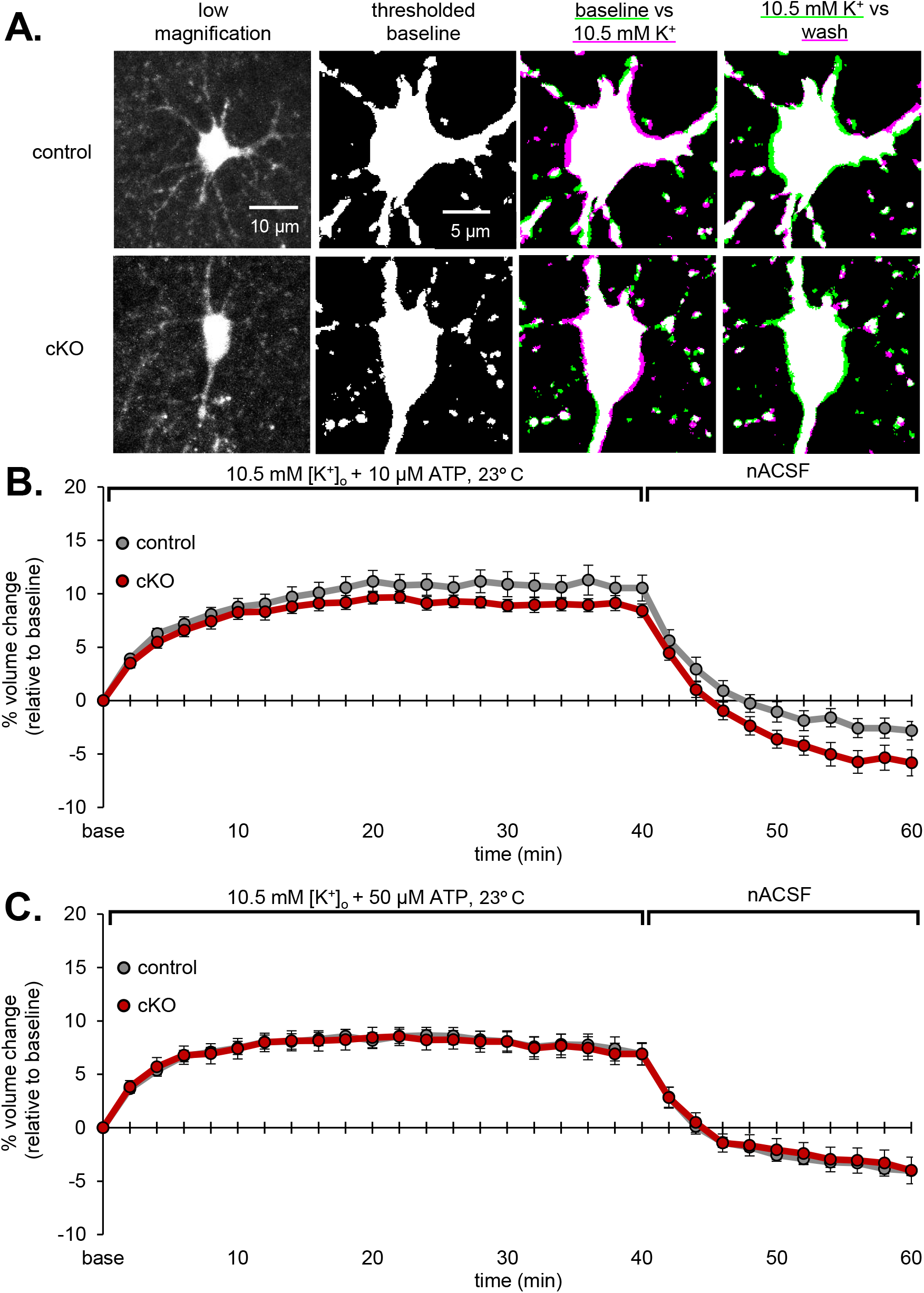
VRAC cKO does not affect the astrocyte swelling response in 10.5 mM [K^+^]_o_. **(A)** (Far Left) Low magnification images of SR101-labeled astrocytes in stratum radiatum of acute hippocampal slices in control (top) and VRAC cKO (bottom) tissue. (Middle Left) Zoomed in and cropped images focus on the astrocyte soma to permit fast z-stack acquisition and optimal analysis of cell volume. (Middle Right) Pseudocolored images were created by merging images from an earlier time point (green) with a later time point (magenta). Magenta regions indicate membrane expansion relative to baseline after 40 minutes in elevated [K^+^]_o_ solution. (Far Right) The end of the wash period is overlaid with the timepoint corresponding to peak volume increase. Green regions indicate where the cell volume has reduced relative to the end of the experimental period, demonstrating volume recovery. **(B)** Baseline volume is established in nACSF and set to 0%. Each subsequent time point is expressed as a percent change relative to baseline such that a value above baseline represents swelling and a value below baseline represents shrinking. Each line represents the average volume response of several astrocytes, and error bars represent standard error of the mean. VRAC intact (gray, n = 12 cells) and VRAC cKO (red, n = 14 cells) astrocytes display roughly the same volume response to 10.5 mM [K^+^]_o_ supplemented with 10 µM ATP. **(C)** Similarly, control (gray, n = 13) and VRAC cKO (red, n = 10) astrocytes exhibited the same amount of swelling in 10.5 mM [K^+^]_o_ supplemented with 50 µM ATP. Note lack of RVD in these conditions for either genotype.

Consistent with our past findings (Walch et al., 2020, 2022), exposure to 10.5 mM [K^+^]_o_ triggered robust and rapid swelling in control astrocytes (**Fig. 2B**). On average, control astrocytes reached a maximum of ∼11% volume increase in 10.5 mM [K^+^]_o_, while VRAC cKO astrocytes swelled to a maximum of ∼9%. Cells reached this volume after ∼18 minutes of exposure to 10.5 mM [K^+^]_o_ then reached a plateau until the end of the application. For both groups, cells began to recover immediately upon removal of the ionic challenge. A two-way mixed ANOVA revealed no interaction between solution exposure and genotype (F(2.61, 720) = 1.09, p = 0.36, η^2^ = 0.04).

Hydrolysis of commonly used ATP salts, like sodium ATP, may be a concern in interpreting contributions of ATP to VRAC activity upon cell swelling in 10.5 mM [K^+^]_o_ (Jackson et al., 1994). Thus, we raised the ATP concentration to 50 µM so that, assuming some hydrolysis by ectonucleotidases, there would still be ample extracellular ATP available for VRAC potentiation after reaching equilibrium in solution. The increased ATP concentration did not influence volume responses in control astrocytes (F(2.98, 660 = 1.92, p = 0.14, η^2^) (**Fig. 2B** and **C**, gray lines). We again observed no differences in swelling profiles of VRAC cKO astrocytes compared to controls with 50 µM ATP (F(2.00, 690) = 0.11, p = 0.90, η^2^ = 0.01) (**Fig. 2C**). Surprisingly, astrocytes did not display any compensatory volume reduction indicative of RVD (H. Pasantes-Morales et al., 1994) during 10.5 mM [K^+^]_o_ exposure, as evidenced by the sustained elevation in astrocyte volume that persisted until the ionic challenge was removed. The slowed rate of swelling partway through the experimental period suggests that astrocytes may modulate their K^+^ and water intake to prevent volume increase beyond a certain threshold, which does not appear to be mediated by VRAC. Overall, these findings indicate that moderate increases in [K^+^]_o_ are not sufficient to trigger VRAC-mediated volume regulation.

### VRAC limits the astrocyte swelling response to 40% hACSF

While astrocyte swelling has been observed in both elevated [K^+^]_o_ conditions and hypoosmolar conditions, there are different underlying mechanisms driving water intake (Murphy et al., 2017; Walch et al., 2020) and, therefore, there may be different contributions of VRAC to swelling responses in each model. We examined the role of VRAC in the long-term swelling response to 40% reduction in extracellular osmolarity. This reduction was achieved by diluting the solution with distilled water to a final osmolarity of 180-183 mOsm (40% hACSF). Ten µM ATP was included to potentiate VRAC, which was expected to exaggerate differences in swelling responses relative to VRAC cKO astrocytes. Representative images were rendered as described above, which showed rapid and sustained swelling during the experimental period (**Fig 3A, middle right**) and recovery at the end of the wash period in normosmolar ACSF (nACSF) (**Fig 3A, middle left**). Average volume increases in this condition far exceeded volume increases observed in 10.5 mM [K^+^]_o_, with control astrocytes swelling to a maximum of ∼24% and VRAC cKO swelling to a maximum of ∼33% (**Fig. 3B**). There was a statistically significant interaction between solution exposure and genotype on astrocyte volume (F(2.78, 780) = 5.19, p = 0.003, η^2^ = 0.17), with control astrocytes swelling at a much lower rate and achieving a lower maximum volume relative to VRAC cKO astrocytes. Since VRAC cKO astrocytes achieved a higher volume by the end of the experimental period, their volume remained elevated during the wash period, although the rate of recovery towards baseline was the same as that of control astrocytes.

**Figure 3:**
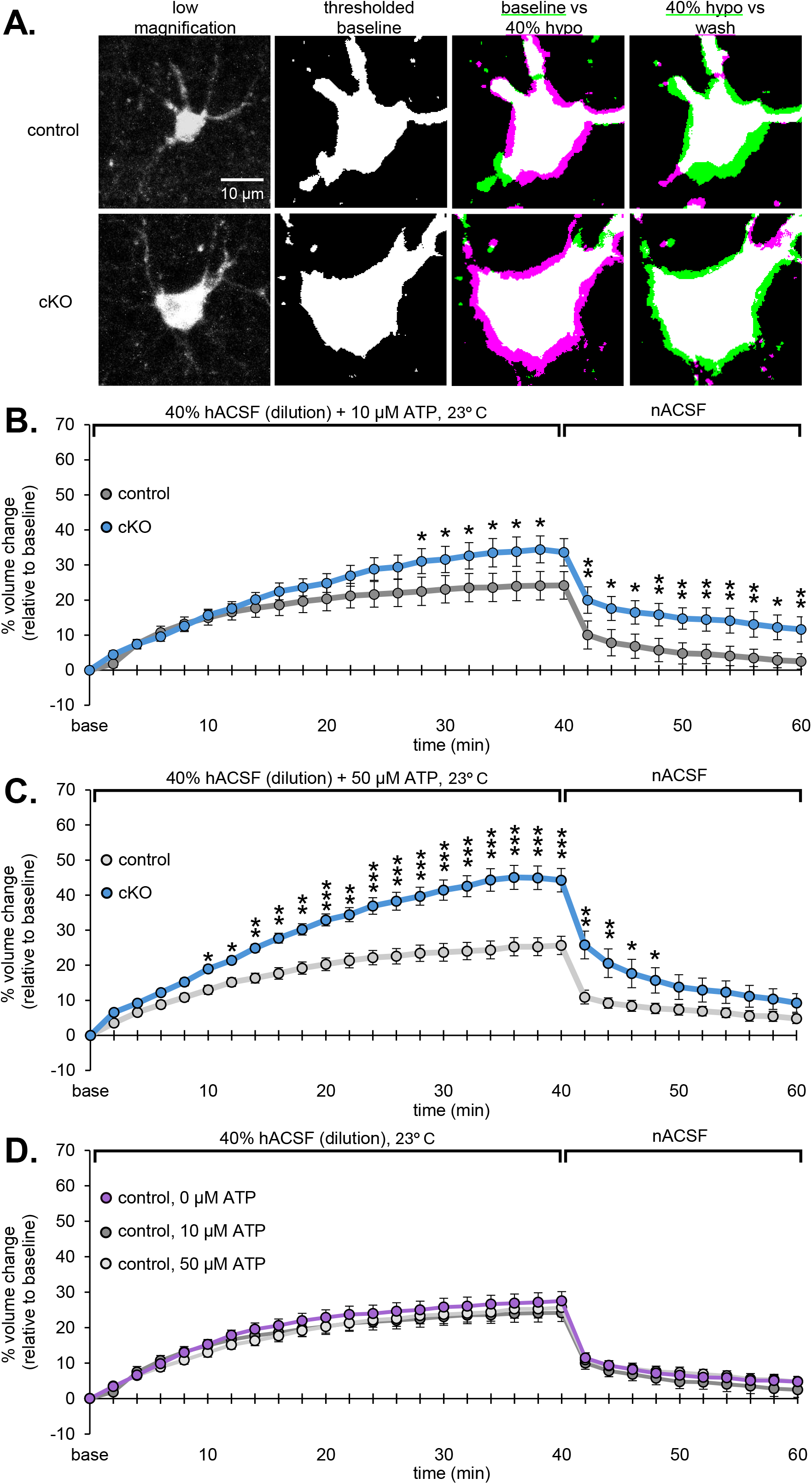
VRAC cKO astrocytes swell more than controls in response to a 40% reduction in solution osmolarity. **(A)** Representative images of VRAC cKO and control astrocytes. Note greater degree of swelling (magenta) in the VRAC cKO astrocyte compared to control. **(B)** On average, VRAC cKO astrocytes (blue, n = 13 cells) swelled significantly more (24% vs. 34%) than control astrocytes after 40 minutes in hACSF (dark gray, n = 15 cells). **(C)** Increasing the ATP concentration to 50 µM led to greater differences in swelling between genotypes, with control astrocytes (light gray, n = 12) swelling significantly less than VRAC cKO cells (26 vs. 44% after 40 minutes; blue, n = 15). Note lack of RVD in either condition. **(D)** Extracellular ATP is thought to potentiate VRAC, possibly leading to greater osmolyte efflux to counter swelling responses. However, ATP had no effect on the amount of astrocyte swelling in control astrocytes (n = 15). Asterisks indicate significant differences between groups as follows: *p<0.05, **p<0.01, ***p<0.001.

Genotype differences in astrocyte swelling persisted in the presence of 50 µM ATP (F(2.40, 750) = 6.24, p = 0.002, η^2^ = 0.20) (**Fig. 3C**). Interestingly, increased ATP had no effect on swelling responses in control astrocytes. However, VRAC cKO astrocytes swelled more in 50 µM ATP compared to 10 µM ATP (**Fig. 3C**, blue lines), suggesting some effect of ATP and/or adenosine on astrocyte swelling responses independent of VRAC. Since our findings in controls contradicted much of the literature regarding the role of extracellular ATP in the activation of VRAC (Kimelberg, 2004; Mongin & Kimelberg, 2005), we further probed whether ATP was required at all for regulation of VRAC in the response to hACSF. We found that VRAC- expressing astrocytes achieved the same maximum volume regardless of ATP concentration (F(4.84, 1170) = 0.64, p = 0.67, η^2^ = 0.03) (**Fig 3D**), suggesting that ATP does not play a role in VRAC-mediated volume regulation.

As was the case in elevated [K^+^]_o_, we did not observe a classical RVD response to 40% hACSF at either concentration of ATP. These trends are consistent with the view that RVD does not occur in intact brain tissue (Andrew et al., 1997), suggesting that osmotic challenge alone is not a sufficient trigger for RVD, even when VRAC is intact. However, the increased rate of swelling in VRAC cKO astrocytes relative to control astrocytes suggested that VRAC plays an important role in limiting the maximal swelling response, a mechanism that appears to be triggered *only* by osmotic, and not ionic, challenge.

### VRAC activity in 10.5 mM [K^+^]_o_ and 40% hACSF is not dependent on temperature

The kinetics of ion and water movement are much faster at physiological temperature, and temperature has been shown to be a highly influential variable on astrocyte swelling, both in primary cultured astrocytes and in brain slices (Andrew et al., 1997; Mola et al., 2016; Pasantes-Morales et al., 1994). Transporter and pump activity is higher at physiological temperature (McCutcheon & Lucke, 1926; Swann, 1983), thus enhancing K^+^ intake through the sodium-potassium pump (NKA) in elevated [K^+^]_o_ (Walch et al., 2020) and increasing likelihood of observing RVD under greater swelling induction. To approximate physiological temperatures, normal and experimental ACSF were preheated before being applied to the slice, while the recording was performed at 36° C.

As expected, astrocytes typically swelled more at 36° C than they did at 23° C in response to experimental ACSF (**Fig. 4A and C**). Control astrocytes swelled to an average maximum of 29% above baseline in elevated [K^+^]_o_ and 34% above baseline in 40% hACSF (**Fig. 4B and D**). As was observed at room temperature, the consequence of VRAC cKO was only observed in 40% hACSF (F(2.42, 840) = 4.04, p = 0.016, η^2^ = 0.13), but not in elevated [K^+^]_o_ (F(2.3, 720) = 1.61, p = 0.21, η^2^ = 0.06) at physiological temperature. Interestingly, the increase in swelling at higher temperatures appeared to diminish the effect of VRAC cKO compared to controls under hypoosmolar conditions. We attribute this to an artifact of greater dilution of cytosolic fluorescent indicator in VRAC cKO astrocytes due to greater swelling responses rather than a true difference in swelling, interfering with thresholding analysis. In other words, without dimming of indicator, we would expect VRAC cKO astrocytes to swell much more compared to controls at higher temperatures. This clipping or leveling-off effect was observed on average when astrocytes reached approximately 45% of baseline volume in hypoosmolar conditions.

**Figure 4:**
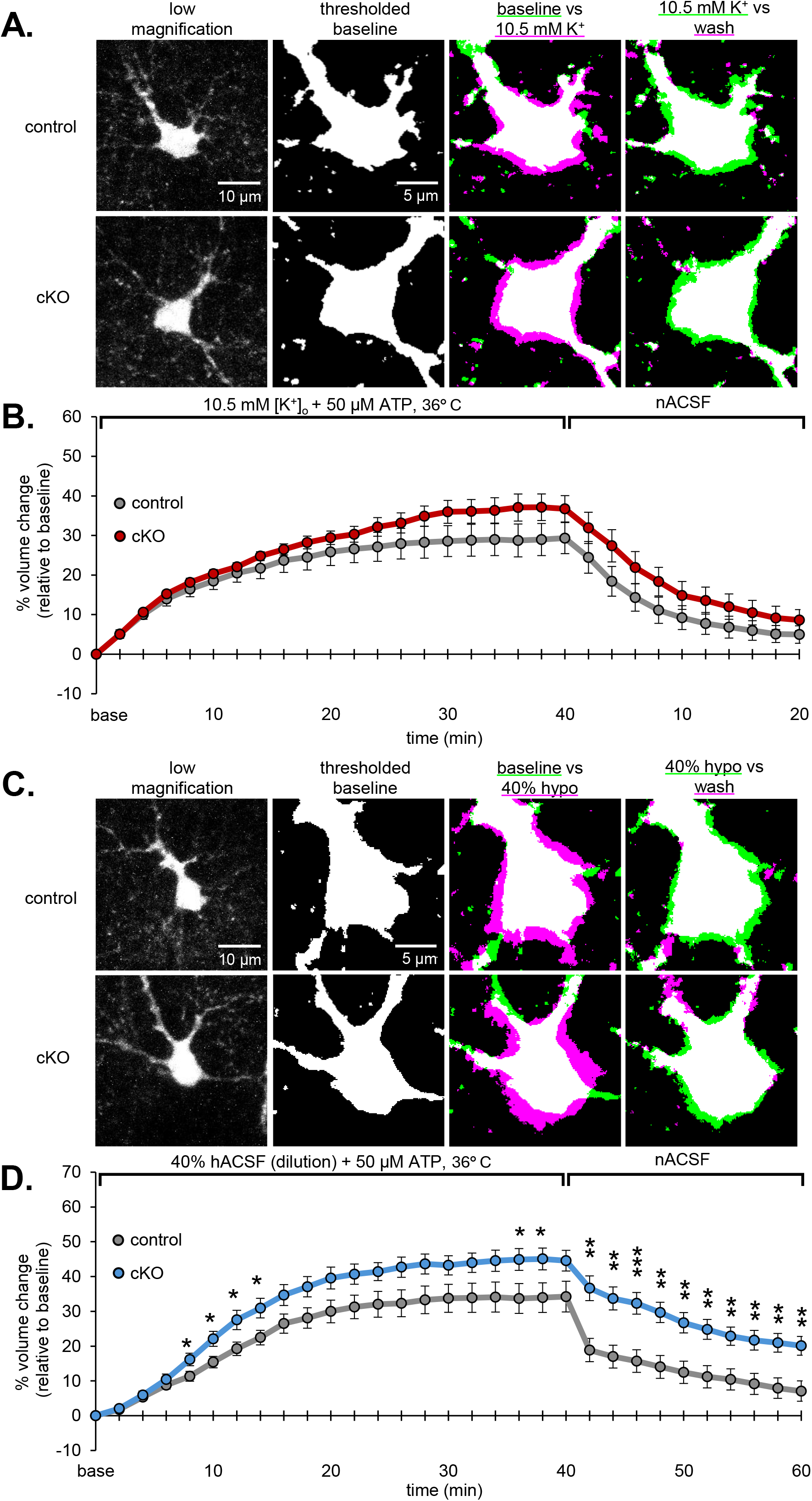
Physiological temperature increases cell swelling but fails to trigger RVD. Volume recordings were performed at 36° C. **(A)** Representative pseudocolor images show expansion (magenta) or reduction (green) of cell volume in 10.5 mM [K^+^]_o_ relative to baseline (middle right) and then to the wash period (far right). **(B)** In general, swelling was greater at physiological temperature than at room temperature, but both control (gray, n = 12) and VRAC cKO (red, n =14) astrocytes respond to 10.5 mM [K^+^]_o_ with the same amount of swelling. **(C)** Representative images of individual VRAC cKO and control astrocytes exhibiting volume responses in hACSF. **(D)** Only hypoosmolar conditions revealed a VRAC cKO effect on astrocyte volume on average. Significant differences in volume between control and VRAC cKO astrocytes are indicated by: *p<0.05, **p<0.01, ***p<0.001.

Taken together, our findings across different ATP doses and temperatures suggest that cells are limited to a certain amount of volume increase that is exceeded when VRAC is ablated, perhaps making them more vulnerable to excessive swelling and, possibly, cell lysis. We did observe that among astrocytes that did not recover upon return to normal ACSF and were thus determined to have lysed, a significant majority were VRAC-ablated cells (control: n = 2; VRAC cKO: n = 6).

### Activation of VRAC is unrelated to the degree of swelling, or to Cl^-^/K^+^ cotransport

Considering the context-dependency of VRAC in different swelling models, we next asked if VRAC participation was related to the amount of swelling in each condition; i.e. the degree of membrane stretch triggered by each model. [K^+^]_o_ levels have been reported to drastically increase past 10.5 mM during extreme pathological conditions such as spreading depression (Kofuji & Newman, 2004; Rutledge et al., 1998; Vyskocil et al., 1972) . Since 10.5 mM [K^+^]_o_ elicited less swelling than 40% hypoosmolar solution even at physiological temperature, we selected a much more dramatic increase of [K^+^]_o_ to match or exceed the level of swelling induced by 40% hACSF. Our previous findings have shown that astrocyte swelling after 5 minutes of exposure to 26 mM [K^+^]_o_ is more than double that in 10.5 mM [K^+^]_o_, and remains restricted to astrocytes (Walch et al., 2020). In the present study, even though swelling in 26 mM [K^+^]_o_ far surpassed swelling in 40% hypoosmolar conditions at physiological temperature, reaching a maximum of 49% in controls and 55% in VRAC cKO, absence of VRAC had no effect on the final astrocyte volume, nor on volume recovery (F(2.99, 690) = 1.18, p = 0.33, η^2^ = 0.05) (**Fig. 5A**). Additionally, 26 mM [K^+^]_o_ elicited the greatest amount of swelling across all conditions, yet was still insufficient to trigger RVD, ruling out the possibility VRAC activation and RVD occurs proportionally to the amount of membrane stretch. Upon comparing peak astrocyte volumes across room and physiological temperatures, it became apparent that VRAC is activated only when external osmolarity is reduced (**Fig. 5B**). These findings support previous work indicating that dilution of intracellular ionic strength is the primary trigger for VRAC activation in astrocytes (Sabirov et al., 2000).

**Figure 5:**
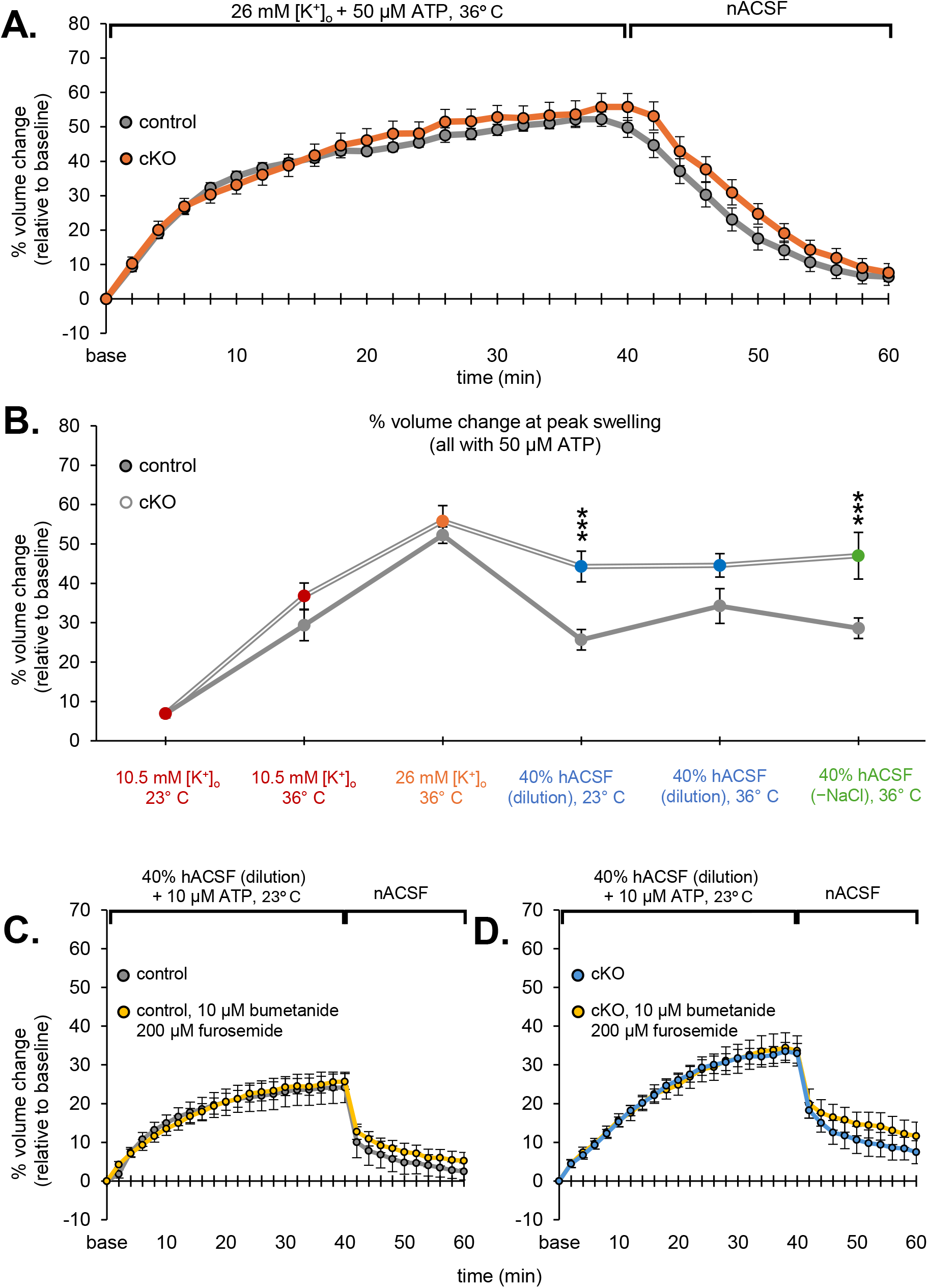
Inhibition of NKCC/KCC does not alter volume differences between control and VRAC cKO astrocytes in hypoosmolar conditions. **(A)** [K^+^]_o_ was raised to 26 mM to generate similar magnitude swelling responses to those observed in 40% hypoosmolar ACSF. Despite increased swelling, the volume response was consistent across both control (gray, n = 12) and VRAC cKO astrocytes (orange, n = 13). **(B)** Astrocyte volume plotted after 40 minutes of exposure to experimental ACSF across all conditions revealed that VRAC regulates astrocyte volume only in conditions where extracellular osmolarity is reduced. **(C)** Pharmacological inhibitors were applied to assess possible contribution from NKCC and KCC cotransporters. Adding 10 µM bumetanide and 200 µM furosemide to 40% hACSF did not affect swelling either in control (n = 12) **(C)** or cKO (n = 10) **(D)** astrocytes. Data from control and cKO astrocytes without antagonists (Fig. 2D) are reproduced here for comparison (gray/blue traces). Significant differences are denoted by: *p<0.05, **p<0.01, ***p<0.001.

Multiple reports have suggested that chloride and potassium movement mediate astrocyte volume changes in various contexts, including cell development and tumor cell migration (Mongin & Orlov, 2001; Pasantes-Morales et al., 1994; Turner & Sontheimer, 2014; Vitarella et al., 1994). Our two swelling models differ in the specific alterations to ACSF ionic composition, affecting the forces driving water entry into the cell. Osmotic strength and [Cl^-^]_o_ remain constant in elevated [K^+^]_o_ ACSF, and outward driving force for Cl^-^ is reduced relative to unswollen cells in nACSF. In contrast, both [Cl^-^]_o_ and [K^+^]_o_ are diluted when extracellular osmolarity is reduced, and outward driving force for Cl^-^ is increased. We therefore asked whether the modified ionic concentration in [K^+^]_o_ creates a condition that is osmotically unfavorable for K^+^ and Cl^-^ efflux, potentially explaining the lack of VRAC participation in these conditions. To test this, we limited routes for Cl^-^ and K^+^ efflux in 40% hACSF conditions by pharmacologically blocking the sodium-potassium-chloride cotransporter (NKCC) and potassium-chloride cotransporter (KCC) with 10 µM bumetanide and 200 µM furosemide (Kahle & Staley, 2008; Macvicar et al., 2002). However, addition of these blockers did not affect the degree of swelling in control or VRAC cKO astrocytes (**Fig 5C, D**), suggesting, again, that VRAC activity is driven by osmolarity of the intracellular compartment, rather than the movement of specific ions across the membrane. Our data also ruled out any potential compensatory activity from other transporters, namely NKCC and KCC.

### Taurine loading is required to elicit RVD

Taurine is an amino acid whose concentrations can range in a location-dependent manner from 1 to 20 µM in the human brain, and from 22 to 89 nM in the rat brain (Palkovits et al., 1986; Ramírez-Guerrero et al., 2022). It is synthesized by a wide variety of cell types, including astrocytes (Vitvitsky et al., 2011). One of the predominant roles assigned to taurine in the brain is osmotic regulation, and several groups have reported on the release of taurine upon astrocyte swelling in both primary astrocyte culture and acute slice preparations (Kimelberg et al., 1990; Kreisman & Olson, 2003; Oja & Saransaari, 2017; Olson & Li, 1997; Vitarella et al., 1994). Taurine uptake is thought to be facilitated by inward sodium transport while taurine efflux is not (Tamai et al., 1995), suggesting that it is unlikely to be transporter mediated (Pasantes-Morales & Schousboe, 1989). Accordingly, taurine efflux has been shown to be inhibited by pharmacological inhibition of anion channels during osmotic challenge (Jackson & Strange, 1993; Pasantes-Morales et al., 1990; Sánchez-Olea et al., 1993). Taurine has also been shown to play an important role in the occurrence of RVD (Kreisman & Olson, 2003). Thus, we set out to test the specific contributions of astrocytic VRAC to taurine release and RVD in intact tissue under different swelling conditions.

Historically, researchers have utilized two methods of reducing osmolarity of experimental ACSF: dilution with distilled H_2_O (Andrew et al., 1997; Cheung et al., 1982; Macvicar et al., 2002; Shao et al., 1994) and reduction of NaCl (Chebabo et al., 1995; Kimelberg et al., 1990; Yang Liu et al., 2023; Mola et al., 2016; Morales-Mulia et al., 2002; Rutledge et al., 1999). We selected 40% hACSF by NaCl removal (−NaCl) for these experiments based on its predilection for eliciting RVD (Kreisman & Olson, 2003), and that reduction of NaCl, even when compensating for osmolarity by the addition of mannitol, dramatically increases taurine efflux in rat cortical astrocytes (Haskew-Layton et al., 2008). At physiological temperature, we found that astrocyte swelling in response to 40% hACSF by dilution (**Fig. 4 C, D**) closely aligned with 40% hACSF by NaCl removal (**Fig. 6A**). As in 40% hACSF by dilution, VRAC cKO astrocytes swelled more than controls following NaCl removal ((F(2.20, 540) = 3.18, p = 0.048, η^2^ = 0.15)). This allowed us to establish that the method of osmolarity reduction did not significantly affect the behavior of control or VRAC cKO astrocytes.

**Figure 6:**
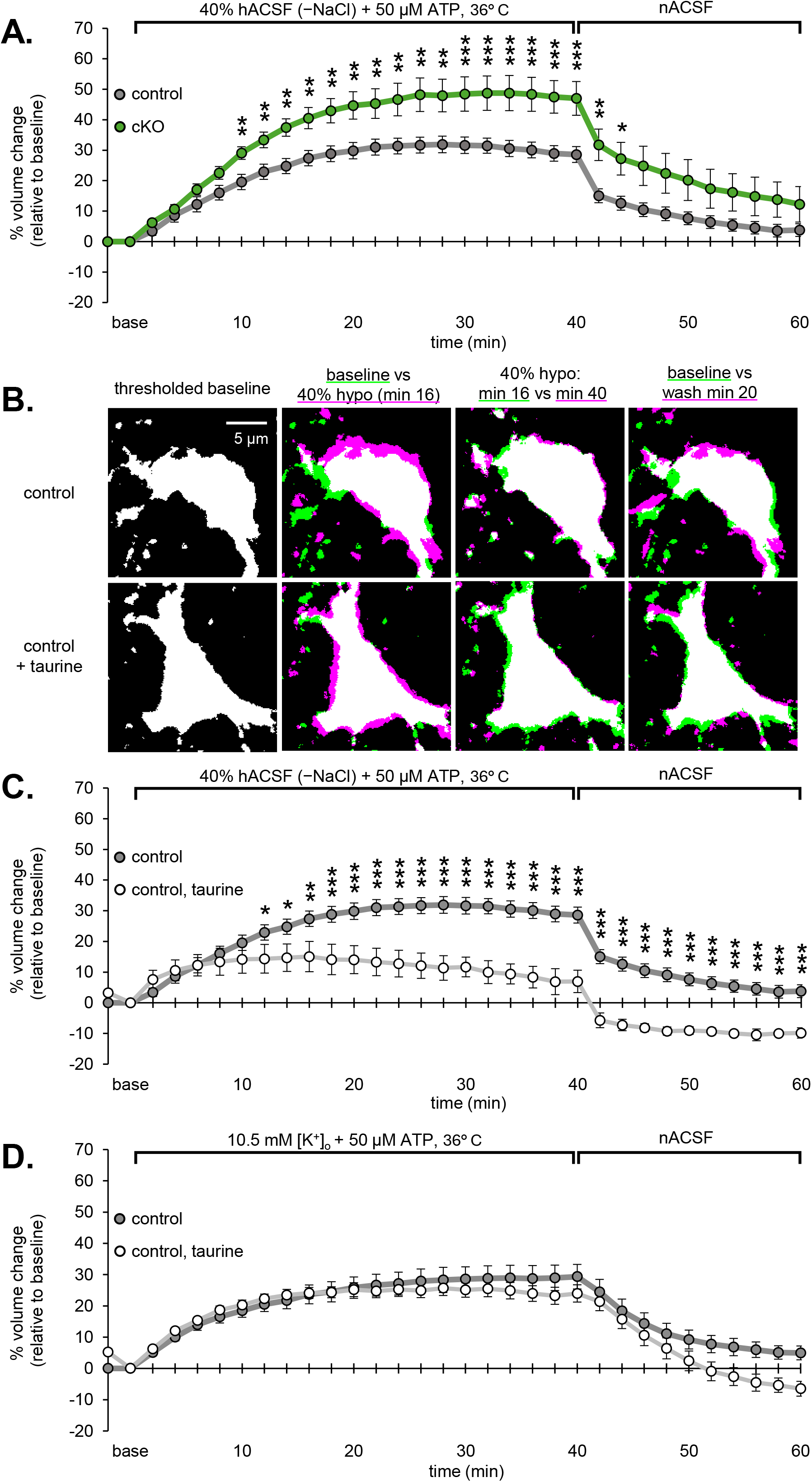
Taurine supplementation results in RVD in hypoosmolar but not elevated [K^+^]_o_ conditions. **(A)** Reducing osmolarity by removing NaCl produced greater swelling responses in VRAC cKO (green, n = 11) compared to control (dark gray, n = 9) astrocytes, similar to findings obtained in hACSF by dilution. There was no RVD in -NaCl hypoosmolar conditions, even at physiological temperature. **(B)** Representative images of control astrocytes before, during, and after exposure to hACSF by NaCl removal with and without taurine. The 16-minute time point was chosen based on when volume differences between taurine-treated vs. untreated cells became apparent. **(C)** In control slices, taurine loading resulted in a peak volume response at 16 minutes, after which time cells began shrinking even in the continued presence of hACSF + taurine for the next 26 minutes. Upon switch back to nACSF, taurine-loaded astrocytes undershot the baseline, a classic hallmark of RVD. Control data without taurine from Fig. 6A is reproduced here (dark gray). **(D)** In 10.5 mM [K^+^]_o_ conditions, taurine supplementation had no effect on the amount of swelling and was insufficient to initiate RVD (light gray, n = 9). Control data without taurine from Fig. 4B are reproduced here. Significance between genotypes indicated by: *p<0.05, **p<0.01, ***p<0.001.

Under physiological conditions, intracellular taurine concentrations are much higher than extracellular concentrations, but taurine levels are significantly depleted during preparation of acute hippocampal slices (Hussy et al., 2000; Law, 1994). To counteract this loss, we supplemented both slicing buffer and ACSF with 1 mM taurine during slice preparation as outlined in Kreisman and Olson (2003), with a minimum exposure time of 90 minutes, which restores intracellular taurine levels closer to physiological conditions. To examine any potential effects of taurine loading itself on astrocyte volume, we obtained a baseline measurement before and after a 30-minute taurine washout period. Astrocytes started at a slightly higher volume immediately following taurine loading. Once baseline volume was re-established in nACSF, we proceeded with application of 40% hACSF and nACSF wash. In non-taurine supplemented controls, swelling proceeded to the end of the experimental period (**Fig. 6B middle right, top**) and resolved at the end of the wash. In contrast, control astrocytes that had been exposed to taurine began to shrink spontaneously after ∼16 minutes (**Fig 6B middle right, bottom**). By the end of the 40% hACSF application, only taurine-loaded astrocytes had decreased their volume from ∼15% to ∼7%, indicating that RVD did occur under these conditions (F(2.57, 450) = 17.47, p < 0.001, η^2^ = 0.54) (**Fig. 6C**). RVD persisted during the wash period (**Fig 6B far right, bottom)** suggesting water efflux was still occurring despite the restoration of normal tonicity, a classic hallmark of RVD (Kreisman & Olson, 2003). Interestingly, taurine loading did not elicit RVD or affect the overall swelling response in 10.5 mM [K^+^]_o_ (**Fig. 6D**), although the rate of recovery was slightly heightened during the wash period. A two-way mixed ANOVA revealed an interaction between taurine loading and solution exposure (F(2.33, 600) = 4.36, p = 0.014, η^2^ = 0.18) but no main effect, suggesting that this difference was minor.

### VRAC is required for RVD

Qualitatively, a stereotypical RVD is characterized by an initial volume increase triggered by a sudden ionic or osmotic challenge, followed by an attempt to reverse swelling and return to baseline conditions (H. Pasantes-Morales et al., 1994). This results in an “undershoot” of the original baseline volume as the challenge is alleviated, indicative of compensatory mechanisms that remain active long after the threat has been removed (Andrew et al., 1997; Kreisman & Olson, 2003).

Having established that RVD only occurs with sufficient intracellular taurine, we then sought to establish a role for astrocytic VRAC in taurine-mediated RVD. The slightly elevated astrocyte volume immediately following taurine loading was consistent between control and cKO astrocytes, suggesting that knockout of VRAC did not interfere with taurine intake. As expected, taurine-loaded astrocytes swelled the same amount in 10.5 mM [K^+^]_o_, regardless of VRAC expression (F(2.42, 558) = 4.36, p = 0.43, η^2^ = 0.047) (**Fig. 7B**). This suggested that neither taurine nor the presence of functional VRAC were sufficient to elicit RVD in 10.5 mM [K^+^]_o_ conditions. In contrast, while taurine-loaded control astrocytes exhibited RVD, taurine- loaded VRAC cKO astrocytes responded to hACSF without observable compensatory volume change (F(2.81, 450) = 12.31, p < 0.001, η^2^ = 0.45) (**Fig. 7C**). This led us to conclude that both VRAC and sufficient intracellular taurine concentrations are required to elicit RVD in 40% hACSF conditions.

**Figure 7:**
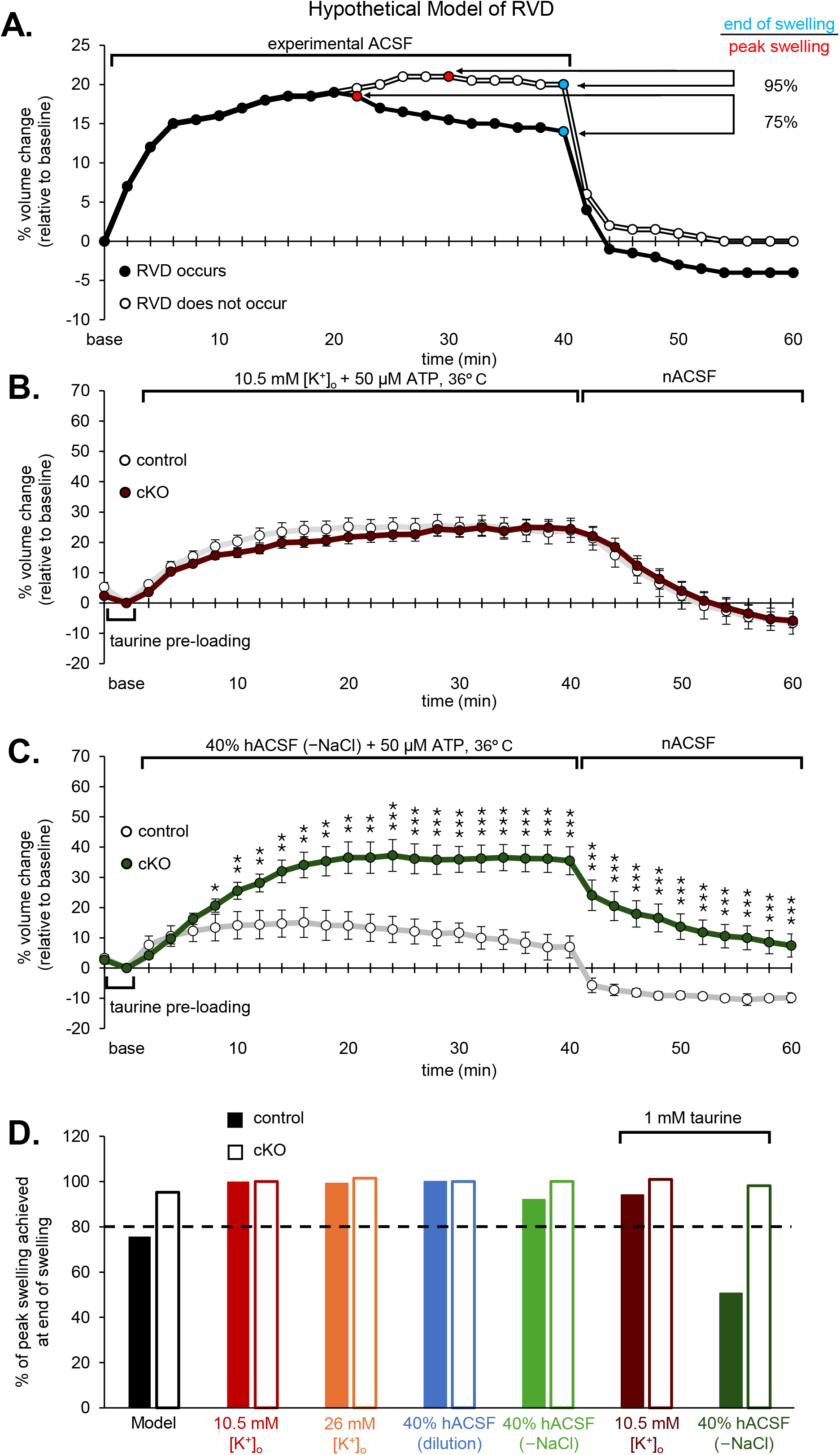
RVD is absent in VRAC cKO astrocytes. **(A)** Hypothetical cellular volume responses to experimental ACSF when RVD does (solid line) or does not (open line) occur. RVD was quantified based on the percent volume change from baseline at the end of swelling (light blue time point) relative to the maximum swelling achieved (red time point). RVD was deemed to have occurred if this ratio was lower than 80% (in other words, compensatory volume decrease exceeded 20%). **(B)** Taurine loading had no effect in either control (light gray, n = 10) or cKO (dark red, n = 10) astrocytes in 10.5 mM [K^+^]_o_. Taurine-loaded control astrocytes are reproduced from Fig. 6D. **(C)** VRAC cKO (dark green, n = 9) abolished the RVD response in astrocytes that occurred in hACSF when taurine was present (light gray, n = 8). Taurine-loaded control traces are reproduced from Fig. 6C. Significant differences are indicated by asterisks as follows: *p<0.05, **p<0.01, ***p<0.001. **(D)** Summary of findings, with the dotted line representing the threshold of recovery required for RVD. When comparing astrocyte volume responses across all recordings performed at physiological temperature in 50 µM ATP, the only condition where RVD was observed was in 40% hACSF by NaCl removal with taurine present (dark green solid bar). In these conditions, RVD was absent in VRAC cKO astrocytes.

**Figure 7D** provides a summary of swelling responses across the different conditions used in our study. If volume reduction occurred that was 80% or more from peak swelling during continued osmotic or ionic challenge (**Fig. 7D, dotted line**), and volume then undershot the baseline upon return to control ACSF, RVD was determined to have occurred. The only conditions where both aspects of RVD – recovery during experimental ACSF and undershoot during the wash period – were observed was in 40% hACSF (–NaCl), at 36° C, with 50 µM ATP, and when VRAC was present (**Fig. 7D, dark green solid bar**).

## Discussion

In this study, we provide compelling evidence that the volume regulated anion channel, VRAC, is required for regulatory volume decrease in intact brain tissue. The occurrence of RVD and the role of VRAC in it are both strongly condition-dependent. Although elevating [K^+^]_o_ and reducing extracellular osmolarity both trigger robust and prolonged astrocyte swelling, VRAC participation in astrocyte volume regulation is limited to hypoosmolar conditions. Administration of taurine was also essential to elicit RVD in hypoosmolar conditions. Physiological temperature combined with reduction of solution osmolarity by NaCl removal were insufficient to induce RVD without taurine present. Overall, many of our findings contradict what has been observed in early reports of RVD and swelling induced osmolyte efflux, both with respect to experimental conditions and time course (H. Pasantes-Morales et al., 1994). These discrepancies may be explained, at least in part, by the differences between *ex-vivo* and cultured astrocyte preparations, where much of the data has been obtained. By performing real-time volume measurements in acute hippocampal slices, and with selective targeting of astrocytic VRAC using transgenic approaches, we were able to investigate the mechanisms underlying long-term astrocyte swelling responses in the intact brain slice preparation. Overall, our findings provide valuable new information on the requirements for astrocyte volume regulation in intact brain tissue and the role of VRAC in these processes.

### Context-dependent mechanisms of astrocyte swelling

While several groups have highlighted the pathological consequences of prolonged reductions in extracellular osmolarity (Hussy et al., 2000; Kilb et al., 2006; Lauderdale et al., 2015) and elevated [K^+^]_o_ (Neprasova et al., 2007; Traynelis & Dingledine, 1988; Walch et al., 2022), the differential mechanisms underlying astrocyte swelling in these two models are rarely compared directly. While neurons readily swell and shrink in responses to changes in solution osmolarity (Murphy et al., 2017; Walch et al., 2022), they are resistant to swelling in elevated [K^+^]_o_ conditions, likely due to pump isoform expression (Walch et al., 2020). Several astrocytic channels have been implicated in water accumulation upon outside K^+^ elevations, including the inwardly rectifying potassium channel (Kir4.1), aquaporin-4 (AQP4), and the sodium-potassium-chloride cotransporter (NKCC) (Binder et al., 2006; Larsen et al., 2014). Our lab has probed several of these and found that water entry into astrocytes is specifically attenuated by inhibition of the sodium-potassium pump (Walch et al., 2020), while Kir4.1, AQP4, the sodium-bicarbonate transporter (NCBe1), or NKCC played no obvious role (Walch et al., 2020). In this study, we also rule out contributions of VRAC to both astrocyte swelling and recovery under high [K^+^]_o_ conditions (**Fig 2B and C, 4B**). Even more drastic increases in [K^+^]_o_ were insufficient to reveal a role for VRAC in astrocyte volume regulation (**Fig 5A**). Although our findings conflict with older reports of astrocyte volume regulation in elevated [K^+^]_o_ in cultured cells (H. Pasantes-Morales et al., 1994; Rutledge et al., 1999), our results are more in line with recent findings obtained from acute brain slices which showed that RVD does not occur in 50 mM [K^+^]_o_, and that pharmacological inhibition of VRAC does not participate in the somatic volume response (Kolenicova et al., 2020).

In contrast to elevated [K^+^]_o_ conditions, swelling in hypoosmotic conditions may be directly tied to VRAC activation, as cells deficient in VRAC swelled significantly more than VRAC-expressing control astrocytes across a range of conditions (**Fig. 3B and C, 4D, 5C-D, 6A**). A few theories have been proposed regarding the mechanisms for water entry into cells under hypoosmolar conditions, including passive diffusion across the membrane, cotransport with other ions, or through AQP4 (Jentsch, 2016; Lauderdale et al., 2015; MacAulay, 2021; Walch & Fiacco, 2022). Our lab has reported that, in response to a 40% reduction in extracellular osmolarity, AQP4 knockout produces a similarly elevated astrocyte volume response relative to wild-type cells (Murphy et al., 2017). It has been recently reported that AQP4 and VRAC play a dual role in RVD in rat hypothalamus, and interact to regulate the effects of hypoosmotic challenge on neuronal activity (Yang Liu et al., 2023). The co-expression or co-activation of VRAC with other volume sensitive channels during osmotic challenge are likely condition- and/or brain region-dependent.

### Mechanisms of VRAC activation

Increased membrane tension as a consequence of membrane stretch had been considered as a possible stimulus for activation of swelling-mediated osmolyte efflux outside of the brain (Browe & Baumgarten, 2003; Hagiwara et al., 1992; Morishima et al., 2000). Additionally, the osmotically activated astrocytic Cl^-^ current has been shown to be diminished upon impedance of actin, which experiences morphological changes upon swelling (Lascola & Kraig, 1996). However, this view has since been challenged on the basis that many cells have surplus membrane which forms slight invaginations in isotonic conditions, minimizing membrane stretch forces when the cell swells (Bertelli et al., 2021; Mongin & Orlov, 2001). In the present study, the greatest degree of swelling occurred in 26 mM [K^+^]_o_, yet this was insufficient to initiate volume regulation (**Fig 5A**). Only reductions in external osmolarity, even when eliciting lower peak swelling responses, appeared to activate VRAC.

It has been proposed that reduced ionic strength, rather than swelling itself, is a potent trigger for VRAC activation (Law, 1994; Yang Liu et al., 2023; Sabirov et al., 2000). Ionic strength may be manipulated independently from osmolarity by reducing concentrations of the main ionic species within ACSF, and compensating for the loss in osmolarity by adding non-membrane permeable solutes such as mannitol (Nilius et al., 1998). This activates a current that is highly similar to the Cl^-^ current evoked upon membrane swelling during exposure to a hypotonic medium (Nilius et al., 1998). Under this theory, cell swelling can be considered an independent process from VRAC activation, and is not required for the initiation of the swelling-induced anion current (Sabirov et al., 2000; Voets et al., 1999). Since water entry into the cell during hypoosmolar challenge is not coupled to the movement of ions, it results in sudden dilution of intracellular ionic strength, something that does not occur when elevating [K^+^]_o_. The fact that VRAC cKO had no effect in any high [K^+^]_o_ conditions further supports the theory that reductions in intracellular ionic strength are critical for activation of VRAC.

The role of Ca^2+^ in astrocytic volume regulation and swelling-induced osmolyte efflux has been a matter of debate. Several different groups have shown both Ca^2+^-dependent and independent signaling pathways associated with activation of VRAC (Akita et al., 2011; Lascola & Kraig, 1996; Yani Liu et al., 2019; Morales-Mulia et al., 2002; Rudkouskaya et al., 2008). Much of the evidence pointing to a role for intracellular Ca^2+^ arises from the importance of ATP on VRAC-mediated organic osmolyte efflux and RVD in cultured cells (Hyzinski-García et al., 2014; Jackson et al., 1994). It has been observed that depletion of extracellular ATP reduces taurine efflux, which is in line with the finding that ATP is critical for anionic current observed during RVD in astrocytes (Jackson et al., 1994; Mongin & Kimelberg, 2005). In the present study, we were surprised to find that augmentation of extracellular ATP up to 50 µM had little to no effect on swelling of control astrocytes, suggesting that ATP activation of purinergic (and/or) adenosine receptors and resulting calcium fluctuations play a minimal role in potentiating VRAC. Surprisingly, we found that VRAC cKO astrocytes actually swelled more in the presence of ATP, suggesting that ATP may exacerbate swelling when VRAC is not present rather than directly potentiating VRAC. One possibility is that ATP potentiation of another channel tied to water entry stimulates isometric volume regulation by VRAC, a process that would be absent in VRAC cKO astrocytes. This highlights the need for further research on potential interactions between VRAC and other volume-sensitive cellular processes, as well as potential consequences of VRAC cKO on other channels and transporters.

The physiological source of ATP also remains under investigation. Swelling-induced ATP release has been reported in cultured hepatocytes (Y. Wang et al., 1996), but there are conflicting reports regarding autocrine ATP signaling by astrocytes (Darby et al., 2003; Mongin & Orlov, 2001). Some reports have also cited the importance of sufficient intracellular ATP in activating VRAC (Jackson et al., 1994; Rutledge et al., 1999; Wilson et al., 2019). The exact ways in which intracellular ATP might impact activation of VRAC are still unknown. Further experimentation would be needed to elucidate the specific mechanisms underlying the role of extracellular as opposed to cytosolic ATP in VRAC activation.

### VRAC as an osmolyte efflux route

The differential mechanisms of swelling in each swelling model provide possible explanations for involvement of VRAC, but not the other model. Volume-sensitive efflux of solutes has been directly associated with Cl^-^ efflux (Hyzinski-García et al., 2014; H. Pasantes-Morales et al., 1990). VRAC is an anion channel, with the primary ionic species being Cl^-^ (Hyzinski-García et al., 2014; H. Pasantes-Morales et al., 1990). External chloride concentration is reduced in both hypoosmolar formulations, but not in elevated [K^+^]_o_. Thus, the outward driving force for Cl^-^ is increased in hypoosmolar solutions but decreased in elevated [K^+^]_o_ (by addition of KCl). This may contribute, at least in part, to the perceived lack of VRAC participation in elevated [K^+^]_o_.

Release of taurine, an organic osmolyte, is another commonly reported consequence of astrocyte swelling (Haskew-Layton et al., 2008; Olson & Li, 1997; H. Pasantes-Morales & Schousboe, 1989). Physiological roles for taurine have been explored in the context of direct RVD mechanisms, as well as a number of cellular processes both within and outside the brain (Ramírez-Guerrero et al., 2022). Reduced taurine levels have been proposed as a potential reason for impaired volume regulatory functions in astrocytes impacted by OGD (Benesova et al., 2009). Surprisingly, although taurine efflux has been shown to occur in both hypoosmolar and elevated [K^+^]_o_ solutions (H. Pasantes-Morales & Schousboe, 1989; Rutledge et al., 1999), our data suggest that VRAC does not participate in regulatory volume function in elevated [K^+^]_o_, with or without taurine. While not considered to be a classical neurotransmitter, taurine has similar structure to GABA and may activate GABA_A_ and glycine receptors (Song et al., 2012; J. Wu et al., 2008). Therefore, it may be a cellular adaptation to combat excitability increases associated with cell swelling. It remains to be seen whether RVD is made possible by taurine efflux specifically, or if efflux of comparable amounts of any other VRAC-permeable solute might have similar effects. Repeating hypoosmolar experiments following loading with other VRAC-permeable osmolytes may address this gap. Pre-loading astrocytes with aspartate or lactate before attempting to evoke swelling and RVD may reveal more about the functional role of taurine efflux in these specific conditions. It may also be useful to determine if the efflux of any solute, VRAC-mediated or otherwise, would be sufficient to trigger RVD. Comparing our findings with taurine or other osmolytes to loading with non-VRAC permeable osmolytes, like sodium bicarbonate (Florence et al., 2012), may address this question.

### Pathological Implications of VRAC Dysregulation

Maintenance of volume is a critical aspect of cellular function, including proliferation, migration, and death (L. Chen et al., 2019; Okada et al., 2020; Pasantes-Morales, 2016). Interestingly, constitutive VRAC knockout from astrocytes did not confer early lethality or physiological abnormalities. However, its deletion from a broad population of neuronal and glial cells resulted in seizures and premature death during early adolescence (Wilson et al., 2021), suggesting that VRAC expression is crucial for healthy brain development and function. VRAC has been implicated in astrocyte function and dysfunction across a variety of disease states, including glioblastoma (Ernest et al., 2005; Ransom et al., 2001), ischemic stroke (Wilson et al., 2019; Yang et al., 2019), and epilepsy (Ghouli et al., 2023). Upon physical degeneration or damage, astrocytes undergo a process called astrogliosis, or reactive gliosis, during which their homeostatic functions become compromised or altered (Pekny & Nilsson, 2005; Pivonkova & Anderova, 2017; Wetherington et al., 2008). The reprogramming of astrocytes into reactive astrocytes may occur acutely, then persist or worsen over longer time scales. Astrocytic VRAC may contribute to pathology across these conditions through two distinct but related mechanisms –the dysregulation of the astrocytic volume response and the release of neuroactive substances. Our recent work has found persistent upregulation of VRAC expression in reactive astrocytes induced by acute neuroinflammation and in the development of temporal lobe epilepsy (Agnew-Svoboda et al., 2022; Ghouli et al., 2023), which could exacerbate pathological or protective roles of VRAC.

Cell swelling is closely associated with the generation of seizure-like activity (Broberg et al., 2008; Traynelis & Dingledine, 1988). Heightened glutamatergic activity as a result of astrocytic swelling has been reported under both hypoosmolar (Kilb et al., 2006; Lauderdale et al., 2015) and elevated [K+]o conditions (Traynelis & Dingledine, 1989; Walch et al., 2022). It has been proposed that swollen astrocytes themselves are the source of glutamate underlying these patterns of hyperexcitability (Angulo et al., 2004; Carmignoto & Fellin, 2006; Wetherington et al., 2008), but the specific mechanisms are under debate. Much of the early VRAC literature identify it as a significant outlet for astrocytic glutamate, potentially underlying increased neuronal synchrony and hyperexcitability associated with seizure-like activity (Pál, 2024; Walch et al., 2022). Even though VRAC did not influence the amount of astrocyte swelling in 10.5 mM [K+]o in the short or long term, we cannot rule out its possible contribution to astrocytic glutamate release under these conditions.

VRAC is also thought to be involved in tissue pathology in the context of ischemic stroke, which, due to the breakdown of fluid barriers in the brain, can involve severe [K^+^]_o_ elevations, excitotoxicity, water imbalance, and cell swelling (Benesova et al., 2009; Feustel et al., 2004; Kahle et al., 2009; Mongin, 2007). The cellular response to these conditions hinges on the ability to regulate volume in in the face of ionic and osmotic imbalances, thus implicating VRAC in the pathological sequelae following stroke. Pharmacological and transgenic interference with VRAC functionality have been shown to have protective effects in mouse models of ischemic stroke (Yang et al., 2019; H. Zhang et al., 2011; Y. Zhang et al., 2008). It is yet unknown whether chronic swelling combined with other changes associated with astrogliosis (Y.-F. Wang & Parpura, 2016) might alter conditions under which VRAC are activated.

Taurine, on the other hand, may have neuroprotective functions beyond RVD (Ademar et al., 2024; K. Wang et al., 2021; Yeon & Kim, 2010). Some studies have reported that taurine inhibits intracellular Ca^2+^ signaling and glutamate-induced excitotoxicity in cultured neurons (W. Q. Chen et al., 2001; El Idrissi & Trenkner, 2004; J.-Y. Wu et al., 2009), suggesting that taurine efflux may not be limited to volume regulation, but that the extruded taurine itself plays a role in mitigating cellular damage in certain pathological conditions.

Overall, VRAC dysfunction in pathological states appears to be highly context-dependent. VRAC deletion may prove to be neuroprotective in some contexts, either by alleviating glutamate-mediated excitotoxicity (Yang et al., 2019), or by a excitatory amino acid-independent mechanism (Balkaya et al., 2023). In other cases, VRAC-mediated RVD and taurine efflux may be essential in relieving cellular edema and reducing glutamate-mediated excitotoxicity and cell damage in neuronal tissue (Leon et al., 2009). Further research is required to investigate the balance between glutamate and taurine efflux, and what governs the harmful or protective effects of VRAC in various pathological contexts.

## Conclusions and Future Directions

In summary, we tested a variety of experimental conditions to test for a role of VRAC in astrocyte volume regulation using a transgenic approach to selectively ablate VRAC in astrocytes. VRAC function is context-dependent, participating in astrocyte volume regulation in hypoosmolar conditions when intracellular ionic strength is reduced, but not in elevated K+ where astrocyte water entry is coupled to intracellular K+ accumulation. Supplementation with taurine, which is otherwise diminished in brain slice preparations, was required to observe an RVD in response to hACSF in control astrocytes, which was completely abolished by removal of VRAC. Our data provide definitive evidence for a role for VRAC in astrocyte volume regulation and RVD in intact brain tissue, while shedding light on the role of ATP in this process.

LRRC8A and its importance to VRAC function is a relatively recent discovery (Formaggio et al., 2019; Qiu et al., 2014), and investigation into how specific VRAC subunit composition may affect channel conductance and osmolyte efflux properties are only beginning to emerge (Syeda et al., 2016). For example, it is unknown how different heteromeric configurations of VRAC subunits might confer different volume regulatory properties, or whether different VRAC channel subtypes are unevenly distributed in astrocyte soma vs. processes. The vast majority of astrocyte volume is distributed in the fine processes, and it is these processes that associate closest with the brain’s vasculature and synapses. A limitation of our work is the difficulty in detecting SR101-labelled fine processes using non-super resolution imaging techniques. Advanced fluorophore loading and imaging capabilities may make volume measurements in smaller astrocytic compartments possible.

Prolonged astrocyte swelling can be extremely detrimental to overall brain function through a suite of direct and secondary effects (Reed & Blazer-Yost, 2022). The experiments performed here were conducted in naïve tissue, but chronic pathologies or injuries may affect VRAC function and, by extension, volume regulation in reactive astrocytes. For example, recent data suggests that astrocytic VRAC is upregulated in a mouse model of mesial temporal lobe epilepsy, implicating either volume dysregulation or excitatory amino acid release as a potential disease mechanism (Ghouli et al., 2023). Given the context-dependent activation of VRAC in otherwise healthy tissue, we can expect that both the type and severity of tissue insult may also determine the extent of VRAC dysregulation. This may be addressed by studying VRAC expression and functional properties across several different pathological contexts, including disordered water intake, systemic inflammation, and/or physical brain injury.

## Acknowledgements

This work was supported by NINDS: Grant/Award Number 1R01NS136434-01A1. We would like to acknowledge Dr. Rajan Sah for providing us with SWELLfl/fl mice.

